# Exploring dynamic brain oscillations in motor imagery and low-frequency sound

**DOI:** 10.1101/2024.03.26.586734

**Authors:** W. Dupont, B. Poulin-Charronnat, C. Madden-Lombardi, T. Jacquet, P. Pfister, T. Dos Anjos, F. Lebon

## Abstract

While both motor imagery (MI) and low-frequency sound listening have independently demonstrated the ability to modulate brain activity, there remains an unexplored frontier regarding the potential synergistic effects that may arise from their combined application. Any further modulation derived from this combination may be relevant for motor learning and/or rehabilitation. We conducted an experiment probing the electrophysiological activity of brain during these processes. By means of EEG, we recorded alpha and beta band power amplitude, which serve as markers of brain activity. Twenty volunteers were instructed to i) explicitly imagine finger flexion/extension movements in a kinaesthetic modality, ii) listen to low-frequency sounds, iii) imagine finger flexion while listening to low-frequency sounds, and iv) stay at rest. We observed a bimodal distribution, suggesting the presence of variability in brain activity across participants during both MI and low-frequency sound listening. One group of participants (12 individuals) displayed increased alpha power within contralateral sensorimotor and ipsilateral medial parieto-occipital regions during MI. Another group (8 individuals) exhibited a decrease in alpha and beta band power within sensorimotor areas. Interestingly, low-frequency sound listening elicited a similar pattern of brain activity within both groups. Surprisingly, the combination of MI and sound listening did not result in additional changes in alpha and beta power amplitudes compared to these processes in isolation, regardless of group. Altogether, these findings shed significant insight into the brain activity and its variability generated during MI and low-frequency sound listening. Nevertheless, it appears that the simultaneous engagement of MI and low-frequency sound listening could not further modulate alpha power amplitude, possibly due to concurrent cortical activations. This prompts us to inquire whether administering these interventions sequentially could uncover any additional modulation.

## Introduction

Motor imagery (MI) is defined as the explicit mental representation of a movement along with the simulation of its sensorimotor consequences, without any associated motor output (Denis, 1989). The utilization of MI has demonstrated its efficiency in enhancing both motor performance and rehabilitation (Ladda et al., 2021; Malouin et al., 2013; Martin et al., 1999; Ranganathan et al., 2004; Ruffino et al., 2017; Slimani et al., 2016). These benefits may be ascribed, at least partially, to the activation of sensorimotor brain regions that overlap those engaged during actual movement (Jeannerod, 2001; Sharma & Baron, 2013). Indeed, numerous studies employing fMRI have posited that MI engages an extensive frontoparietal network and subcortical structures, including the primary motor cortex, supplementary motor area, parietal lobe, and cerebellum (Habas & Manto, 2014; Hardwick et al., 2018; Hétu et al., 2013; Lotze et al., 1999). Nevertheless, although fMRI offers an outstanding spatial resolution, its limited temporal resolution restricts the ability to probe dynamic and subtle cortical changes, such as those underlying MI.

Thereby, EEG recording, particularly the examination of the alpha (8–12 Hz) and beta bands (13–20 Hz), has also emerged as a valuable method for probing cerebral functioning with an excellent temporal resolution, in both motor and cognitive processes. Some researchers have indicated that MI increased brain activity in contralateral sensorimotor or posterior areas, often characterized by a power decrease or an event-related desynchronization in alpha and beta bands (Brinkman et al., 2014; de Lange et al., 2008; Di Nota et al., 2017; McFarland et al., 2000; Pfurtscheller, 2000; Pfurtscheller & Neuper, 1997; Pfurtscheller et al., 1997; Putzolu et al., 2022). However, other researchers have suggested opposite outcomes in the same areas with a power increase or event-related synchronization (Jacquet et al., 2021; Marks & Isaac, 1995; Neuper et al., 2006) or no modulations at all (Kim et al., 2014). While an increase in alpha and beta power might be interpreted as an active inhibition or a disengagement of cortical areas, a decrease in alpha and beta power is thought to correspond to a heightened level of cortical activity, reflecting activated neural clusters ready or primed for sensory, motor, or cognitive processing. Several researchers have suggested that these discrepancies may be attributed to interindividual variations (Höller et al., 2013; Wriessnegger et al., 2020). They argue that studies attempting to identify a common pattern for all participants without taking into account interindividual variability do not capture the true landscape of brain activity during MI. We will return to this idea in our predictions below.

Another approach for modifying both cortical activity and behavioral performance involves the utilization of low-frequency sound stimulation. This method posits that specific sound frequencies, due to their vibratory properties, can be employed to stimulate the brain through auditory pathways, hence regulating oscillatory brain activity (Bartel et al., 2017). Furthermore, the act of listening to low-frequency sounds appears to enhance the performance of rapid dynamic contractions in healthy individuals (Fernandez-Del-Olmo et al., 2014), and also demonstrates promise in improving motor rehabilitation for individuals with fibromyalgia or spasticity following stroke (Chatain et al., 2018; Naghdi et al., 2015). Hence, it seems that both MI and low-frequency sound listening may influence cortical activities and motor performance. In light of these insights, Dos Anjos et al. (2023) recently investigated whether the combination of MI and low-frequency sound listening could improve the neuromotor rehabilitation of patients following knee injuries. Their key result was an improved muscle activation associated with a reduced motor deficit after rehabilitation using MI and low-frequency sound listening. Nevertheless, the combined use of MI and low-frequency sound listening was not compared to MI or to listening to low-frequency sounds in isolation. Therefore, the physiological evidence currently available remains insufficient to definitively ascertain whether the combined use of MI and low-frequency sound listening can produce more substantial brain activity modulations or behavioral improvements than MI or low-frequency sound listening alone.

The present paper aims to shed new light on this fundamental gap by investigating modulations in alpha and beta band activities during MI, low-frequency sound listening, and the combination of both. Given the idea stated above that certain individuals exhibit an increase, while others experience a decrease or no modulation in alpha and beta band activity during MI (Höller et al., 2013; Wriessnegger et al., 2020), our predictions will take into account interindividual variability. First, for individuals exhibiting an increase in brain activity during either MI or low-frequency sound listening in isolation, we expect to observe a further increase in brain activity (power decrease) during the combination of MI and low-frequency sound listening within sensorimotor and parieto-occipital regions. Second, for individuals showing a decrease or no modulation of brain activity during the combination of MI and low-frequency sound listening, we expect to observe a tendency to increase brain activity (from power increase to power decrease) during MI or low-frequency sound listening in isolation. Through this investigation, our aim is to provide a comprehensive understanding of the neural responses elicited by MI, low-frequency sound listening, and the combination of both.

## 1. Material and method

### 2.1 Participants

Twenty right-handed individuals were included in the study (12 women; M_age_ = 22.5 years old; range: 18–30 years old). Participants’ handedness and imagery vividness were assessed by the Edinburgh inventory (range: 0.33–1; Oldfield 1971) and the revised version of Movement Imagery Questionnaire (range: 35–51, showing good to great imagery vividness, 56 being the highest score possible at the questionnaire; Hall & Martin 1997), respectively. All participants were French native speakers and had normal or corrected-to-normal vision, without neurological, physical, or psychiatric disorders. A local Ethics Committee (CER STAPS, IRB00012476-2023-15-12-284) in accordance with the Declaration of Helsinki (excluding preregistration) approved the experimental protocol and procedures. All participants provided written consent to confirm their participation.

### 2.2 Procedure and stimuli

The participants volunteered to participate in a single experimental session that included EEG and electromyographic (EMG) recordings. These measurements were collected in an acoustically and electrically isolated booth during periods of rest, low-frequency sound listening, kinaesthetic MI, or the combination of MI and low-frequency sound listening.

In the low-frequency sound listening condition, participants listened to low-frequency sounds over headphones played by a low-frequency sound generator (Alphabox® system, Allyane), at frequencies spanning from 200 to 400 Hz, with 50 Hz steps. Each sound lasted approximately 10 s (9.4 ± 0.2 s) and the intertrial interval was between 10 and 13 s.

In the MI condition, participants were instructed to imagine 30 trials of dynamic flexion/extension movements of fingers in a kinaesthetic modality. Each trial lasted approximatively 10 s (9.4 ± 0.2 s) and the participants imagined the flexion/extension movement at a pace of 1 Hz. The intertrial interval was between 10 and 13 s. The following instructions were provided: “Try to imagine yourself flexing and extending your fingers, by feeling the muscle contraction and tension as if you were doing the movement”.

In the combined task (MI + Sound), participants performed the same MI task immediately upon sound onset and stopped imagining when the sound disappeared (sounds presented as in the low-frequency sound listening condition).

### 2.3 EEG recording

EEG activities were continuously recorded through the BioSemi system with 64 electrodes according to the 10–20 International system. Eye blinks and horizontal eye movements were monitored with an electrode placed under the left eye and electrodes positioned at the left and right canthi, respectively. Two additional external electrodes were positioned over the left and right mastoids (A1 and A2). Throughout the recording, the common-mode sense (CMS) and the driven right leg (DRL) electrodes of the BioSemi system functioned as active and passive reference electrodes. Electrophysiological signals were acquired with the ActiView software and digitized at a sampling rate of 512 Hz.

### 2.4 EMG recordings

The EMG signal was acquired using surface electrodes with a 10 mm-diameter (Contrôle Graphique Médical, Brice Comte-Robert, France) placed over the Flexor Digitorum Superficialis (FDS) and Extensor Digitorum Superficialis (EDS) muscles of the right forearm. To minimize interference and noise in the EMG signal (< 20 μV), the skin was carefully shaved and cleaned prior to electrode placement. The EMG signals were then amplified and bandpass filtered on-line (10–500 Hz, Biopac Systems Inc.) and digitized at 2000 Hz for subsequent off-line analysis. Specifically, we quantified the root mean square of the EMG signal (EMG_rms_) from each muscle.

### 1.5 Data analysis

Using G* Power (version 3.1.9.4., Faul et al., 2007), we estimated that 18 participants would be needed, based on a large effect size of 0.33 and a power of 0.8 (Di Nota et al., 2017). Due to potential loss of data (10%), we recruited 20 participants in total.

EEG data preprocessing was conducted via Matlab (The MathWorks, Natick, Massachusetts, USA) and the EEGLAB toolbox (Delorme & Makeig, 2004). The EEG signal was bandpass filtered from 1 to 50 Hz and rereferenced to the average of mastoid electrodes (A1 and A2). Electrodes presenting a noisy signal were visually identified, and their signal was removed and interpolated. The Runica routine was applied to correct eye-movement artifacts through independent component analysis (ICA), and components reflecting ocular artifacts were removed based on visual inspection. EEG signals were segmented into 8-s epochs from 1 s to 9 s after the onset of the low-frequency sound. Noisy epochs were rejected by visual inspection and excluded from further analysis. Then, spectral analysis was performed with Fast Fourier Transform (FFT) using the *spectopo* function of the EEGLAB toolbox for alpha (8–12 Hz) and beta (13–20 Hz) frequencies. To perform analyses on spectral data, two regions of interest (ROIs) were selected a posteriori. The first ROI corresponds to the sensorimotor cortex and includes the electrodes Cz and C3. The second ROI corresponds to the right and central parieto-occipital cortex and encompasses the electrodes POz, Pz, P2, P4, P6, PO4, and PO8.

Statistics and data analyses were performed using the Statistica software (Stat Soft, France). Normality, sphericity, and homogeneity of the data were checked with Shapiro-Wilk, Mauchly and Levene tests, respectively. The data are presented as mean values (± SD), and the alpha value was set at .05. Partial eta squared is reported, and thresholds for small, moderate, and large effects were set at .01, .07, and .14, respectively (Cohen, 1988). For paired *t*-tests, Cohen’s *d* thresholds for small, moderate, and large effects were set at .2, .5, and .8, respectively (Cohen, 1988).

To evaluate the impacts of MI, low-frequency sound listening, and their combined influence, a series of deliberate, a priori comparisons were conducted, guided by our hypotheses (Howell, 2012). To avoid a false negative when comparing the four conditions within one repeated measures ANOVA, we performed a separate hypothesis-driven analysis. Based on the literature showing opposite brain modulations during MI, in addition to a qualitative observation of our data with distribution and correlations analysis, we used a cluster analysis to reveal the existence of a bimodal distribution with two distinct participant cohorts characterized by divergent electrophysiological patterns. Then, we contrasted the alpha and beta power density between MI and rest conditions, shedding some light on the distinctive impact of MI on neural activity. We then examined the effects of listening to low-frequency sounds in comparison to rest. Furthermore, we explored the interplay between MI and low-frequency sound listening by comparing their combined effect against that of MI or low-frequency sound listening in isolation. For each comparison, we performed repeated measures ANOVAs with a Group factor (see below for details).

Finally, to ensure that EEG activity was not biased by muscle activation, we used a Friedmann ANOVA to compare the EMG_rms_ within the FDS and EDS muscles for each condition (See Supplementary section).

## 2. Results

### 2.1 Variability of cortical changes and clustering on motor imagery data

As outlined in the introduction, certain individuals exhibit an increase, while others experience a decrease or no modulation in alpha and beta band activity during MI (Höller et al., 2013; Wriessnegger et al., 2020). To take such interindividual variability into account in all tasks, we qualitatively scrutinized the data distribution. We noticed a distribution of cortical changes, from decrease to increase of alpha power, during MI in comparison to rest, as well as during low-frequency sound listening and the combination of both tasks (Figure 1). To go further, we found linear correlations between the modulation of alpha activity during MI, Sound, and MI + Sound for both sensorimotor and parieto-occipital regions (Figure 1): individuals exhibiting increased power during MI also demonstrated such modulation during Sound and MI + Sound, and conversely [MI and Sound: sensorimotor: *r*(18) = .700, *p* = .001; parieto-occipital: *r*(18) = .602, *p* = .005; MI and MI + Sound: sensorimotor: *r*(18) = .583, *p* = .007; parieto-occipital: *r*(18) = .583, *p* = .007; Sound and MI + Sound: sensorimotor: *r*(18) = .707, *p* < .001; parieto-occipital: *r*(18) = .467, *p* = .038]. These correlations highlight the interindividual variability in brain changes and consistent patterns across tasks.

**Figure 1.**
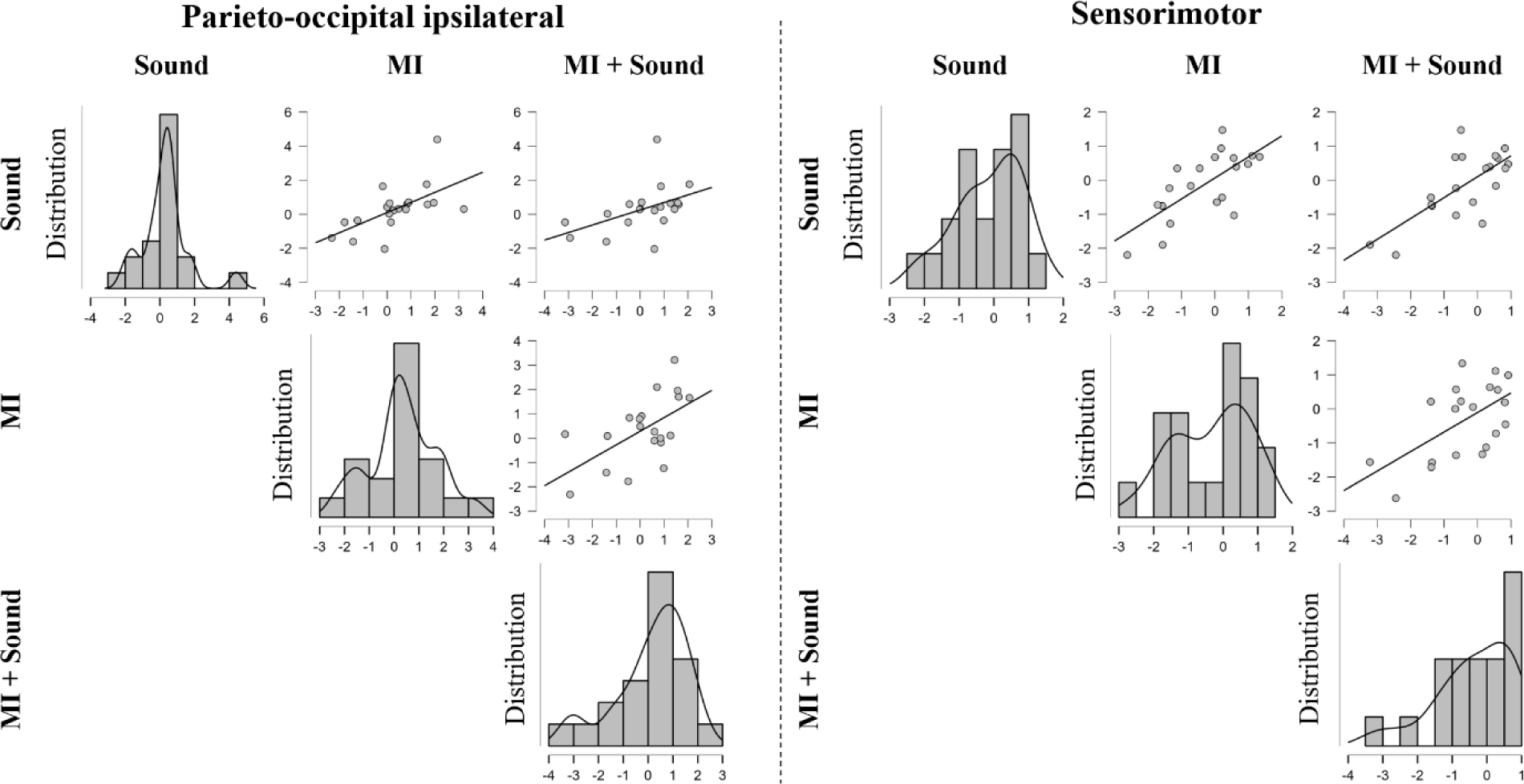
Distribution and correlation analysis of alpha power modulation across tasks and two brain regions of interest. For each plot, data are normalized to rest. MI = Motor imagery, Sound = low-frequency sound listening.

Building upon the literature showing conflicting brain activation during MI and the qualitative observation of our data, we noted the existence of a bimodal distribution with two distinct participant cohorts characterized by divergent EEG patterns. Therefore, we used the silhouette coefficient as a metric to ascertain the optimal quantity of subgroups. The silhouette coefficient plays a pivotal role when selecting the number of subgroups/clusters, thus bolstering the robustness and reliability of the clustering procedure (Rousseeuw, 1987). The silhouette score is a value ranging from 0 to 1, with values closer to 1 indicating a higher optimality of the cluster number. Since MI has demonstrated the ability to influence brain activity at the sensorimotor level, we used the difference between MI and rest in the sensorimotor areas (C3-Cz) to conduct silhouette score analysis. This analysis revealed that our dataset likely comprises two distinct clusters (see Table 2 in supplementary section). Following this, a *K*-means analysis was employed to split the data into two distinct clusters of 12 (Group 1) and 8 participants (Group 2). Then, for each comparison, we performed repeated measures ANOVAs with Group as a between-subject factor.

### 2.2 Motor imagery

For sensorimotor regions (C3-Cz), we observed a main effect of Task for both alpha [*F*(1, 18) = 18.898, *p* < .001, 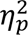 = .512] and beta bands, [*F*(1, 18) = 11.157, *p* = .004, 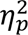 = .383], and a Task by Group interaction for alpha [*F*(1, 18) = 65.608, *p* < .001, 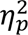 = .784], but not beta band [*F*(1, 18) = .469, *p* = .502, 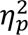 = .025]. Nevertheless, no main effect of Group was observed for either alpha [*F*(1, 18) = .231, *p* < .637, 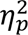 = .013] nor beta band [*F*(1, 18) = .048, *p* = .828, 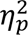 = .003]. As expected from the clustering analysis, using post hoc comparisons with Tukey corrections, we found larger alpha power amplitude during MI (61.66 ± 4.12) in comparison to Rest (61.20 ± 3.87, *p* = .03, *d* = .88) for Group 1, and smaller alpha power amplitudes during MI (59.86 ± 3.37) in comparison to Rest (61.36 ± 3.19, *p* < .001, *d* = 2.74) for Group 2 (see Figure 2). Regardless of group, this was accompanied by a smaller beta power amplitude for MI (55.78 ± 2.53) in comparison to Rest (56.32 ± 2.46, *p* = .004, *d* = .74).

**Figure 2.**
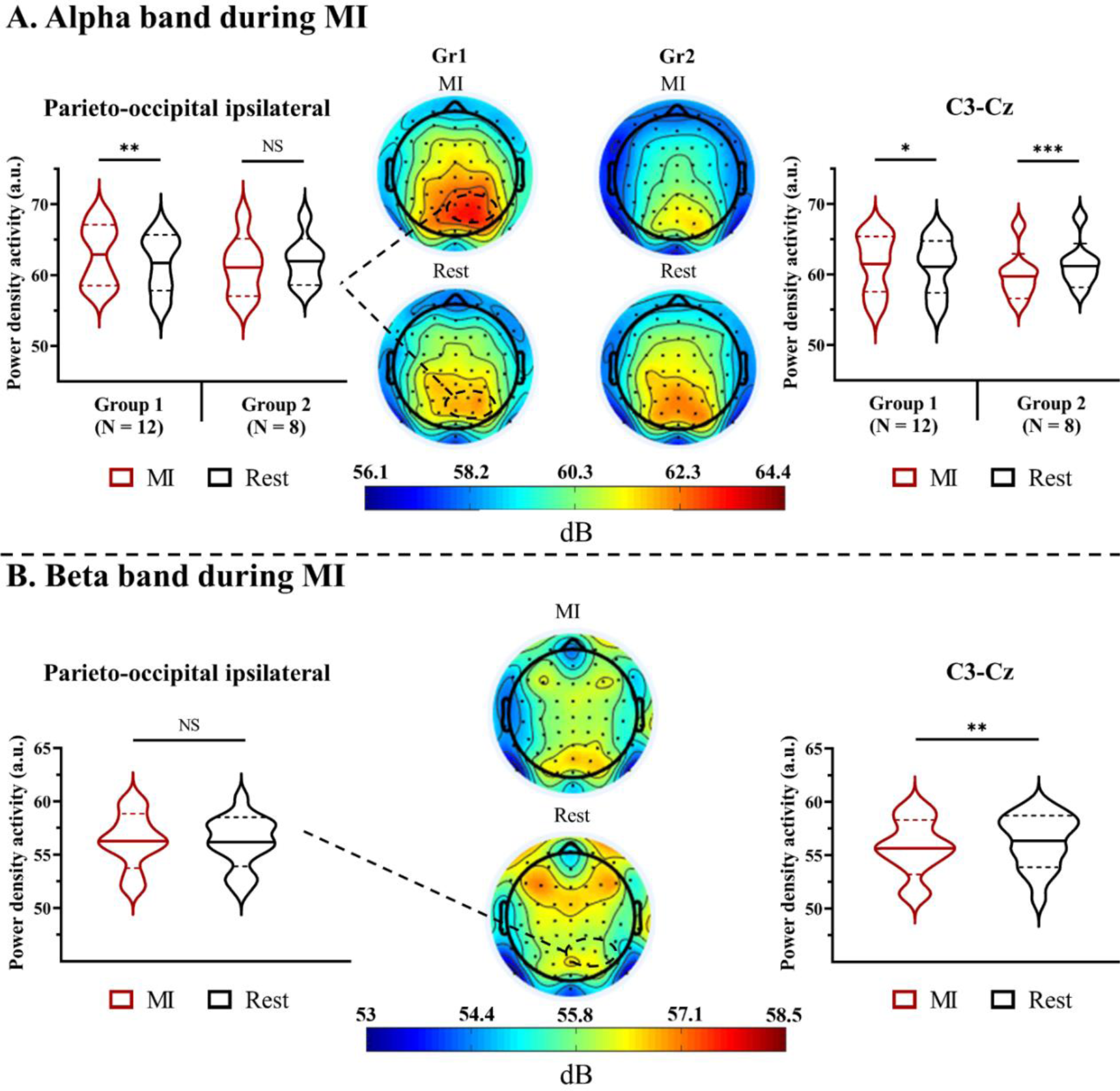
Alpha **(A)** and beta **(B)** band power during motor imagery (MI). The middle figure represents scalp topography of power density activity for the alpha (8–12 Hz) and beta (13–20 Hz) power at rest and during MI. For the right and left figures, violin plots represent power density activity. Thick and thin horizontal lines mark the mean and SD, respectively. Group difference is indicated for the alpha band, while both groups are combined for the beta band. *= *p* < .05, **= *p* < .01, ***= *p* < .001, NS = Nonsignificant.

For parieto-occipital regions, we did not observe any main effect of Task for alpha and beta bands [alpha: *F*(1, 18) = .526, *p* = .477, 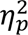 = .028; beta: *F*(1, 18) = .180, *p* = .674, 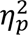 =.010], nor Group [alpha: *F*(1, 18) = .174, *p* = .681, 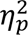 = .009; beta: *F*(1, 18) = .010, *p* = .908, 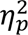 < .001]. However, we observed a Task by Group interaction for alpha band [*F*(1, 18) = 18.544, *p* < .001, 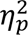 = .507] but not for beta band [*F*(1, 18) = 1.780, *p* = .198, 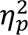 = .090], with a larger alpha power amplitude for MI (62.82 ± 4.25) in comparison to Rest (61.67 ± 3.95, *p* < .005, *d* = 1.14) for Group 1. We did not observe such a difference between MI (61.09 ± 4.06) and Rest (61.90 ± 3.31, *p* = .132, *d* = .82) for Group 2 (see Figure 2). For both regions, the analysis of effect size indicates a moderate to large effect, lending support for our MI findings and their validity for practical application.

### 2.3 Low-frequency sound listening

For sensorimotor regions (C3-Cz), we found no main effect of Task [alpha: *F*(1, 18) = 2.063, *p* = .168, 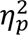 = .103; beta: *F*(1, 18) = .700, *p* = .412, 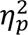 = .038], or Group [alpha: *F*(1, 18) = .076, *p* = .787, 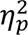 = .004; beta: *F*(1, 18) = .270, *p* = .612, 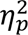 = .015]. However, we observed a Task by Group interaction for both alpha [*F*(1, 18) = 11.415, *p* = .003, 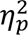 = .388] and beta bands [*F*(1, 18) = 7.300, *p* = .015, 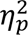 = .288], exhibiting similar alpha power amplitudes for Sound (61.55 ± 3.49) and Rest (61.20 ± 3.87, *p* = .44, *d* = .48) for Group 1, and a smaller alpha power amplitude during Sound (60.50 ± 3.58) compared to that at Rest (61.37 ± 3.19, *p* < .03, *d* = .98) for Group 2 (See Figure 3). Conversely, we observed a marginal larger beta power amplitude for Sound (56.95 ± 2.51) compared to that at Rest (56.38 ± 2.66, *p* = .053, *d* = .71) for Group 1, and no difference between Sound (55.94 ± 2.24) and Rest (56.24 ± 2.31, *p* = .633, *d* = .59) for Group 2.

**Figure 3.**
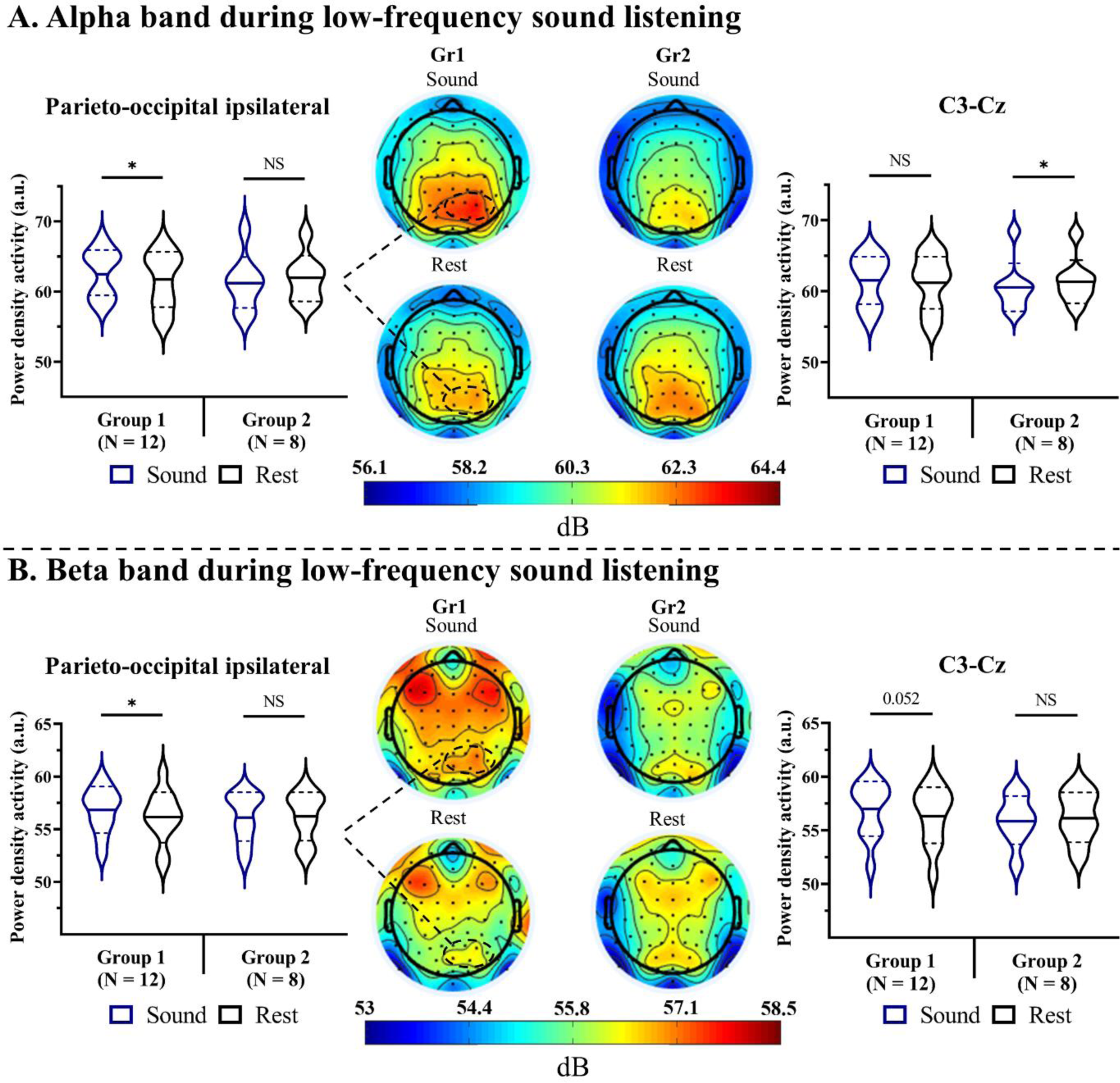
Alpha **(A)** and beta **(B)** band during low-frequency sound listening for Group 1 and Group 2. The middle figure represents scalp topography of power density activity for alpha (8– 12 Hz) and beta (13–20 Hz) power at rest and during sound listening. For the right and left figures, violin plots represent power density activity. Thick and thin horizontal lines mark the mean and SD, respectively. *= *p* < .05.

For parieto-occipital regions, we found no main effect of Task [alpha: *F*(1, 18) = .339, *p* = .568, 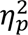 = .018; beta: *F*(1, 18) = 3.390, *p* = .081, 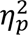 = .159], or Group [alpha: *F*(1, 18) = .090, *p* = .770, 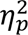 = .005; beta: *F*(1, 18) = .090, *p* = .770, 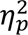 = .004]. However, Task by Group interactions were observed for both alpha [*F*(1, 18) = 11.836, *p* = .003, 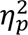 = .397] and beta bands [*F*(1, 18) = 6.220, *p* = .023, 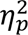 = .257], showing a larger alpha and beta power amplitude during Sound [alpha: 62.67 ± 3.41; beta: 56.81 ± 2.14] compared to that at Rest [alpha: 61.67 ± 3.95, *p* = .024, *d* = .84; beta: 56.16 ± 2.44, *p* = .014, *d* = .89] for Group 1. However, no difference was found between Sound [alpha: 61.20 ± 3.90; beta: 56.13 ± 2.31] and Rest [alpha: 61.90 ± 3.31, *p* = .286, *d* = .79; beta: 56.23 ± 2.29, *p* = .974, *d* = .19] for Group 2 (see Figure 3). For both regions, the analysis of effect size indicates a moderate to large effect, supporting our low-frequency sound listening findings and their validity for practical application.

### 2.4 Combination of motor imagery and low-frequency sound listening

For sensorimotor regions (C3-Cz), we found a main effect of Task for beta band [*F*(1, 18) = 6.826, *p* = .003, 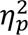 = .275] but not for alpha band [*F*(1, 18) = 1.161, *p* = .324, 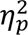 = .060]. However, for both alpha and beta bands, we did not observe any main effect of Group [alpha: *F*(1, 18) = .552, *p* = .467, 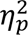 = .030; beta: *F*(1, 18) = .323, *p* = .578, 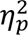 = .018], nor interaction between Task and Group [alpha: *F*(1, 18) = 3.216, *p* = .052, 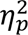 = .151; beta: *F*(1, 18) = 1.111, *p* = .340, 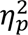 = .059]. Using post hoc comparisons with Tukey corrections, we noticed a smaller beta power amplitude during MI (55.78 ± 2.53, *p* = .003, *d* = .75) or MI + Sound (55.80 ± 2.72, *p* = .004, *d* = .68) than that during Sound (56.54 ± 2.40). Nonetheless, no significant difference between alpha power amplitudes for MI + Sound (60.88 ± 3.62) in comparison to MI (60.90 ± 3.86, *p* = .942, *d* = .06) or Sound (61.13 ± 3.47, *p* = .375, *d* = .31) in isolation were found.

For parieto-occipital regions, for both alpha and beta bands, we found no main effect of Task [alpha: *F*(1, 18) = .216, *p* = .807, 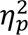 = .011; beta: *F*(1, 18) = .780, *p* = .468, 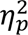 = .041], nor Group [alpha: *F*(1, 18) = .730, *p* = .404, 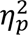 = .039; beta: *F*(1, 18) = .120, *p* = .737, 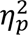 = .006], nor interaction between Task and Group [alpha: *F*(1, 18) = 0.185, *p* = .831, 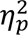 = .010; beta: *F*(1, 18) = 1.460, *p* = .247, 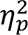 = .075]. Thus, no significant differences emerged for alpha (61.90 ± 4.05) and beta (56.38 ± 2.46) power amplitudes when comparing MI + Sound and MI (alpha: 62.12 ± 4.16, *p* = .745, *d* = .17; beta: 56.29 ± 2.48, *p* = .858, *d* = .15) or Sound in isolation (alpha: 62.07 ± 3.59, *p* = .832, *d* = .12; beta: 56.54 ± 2.18, *p* = .611, *d* = .19; See Figure 4).

**Figure 4.**
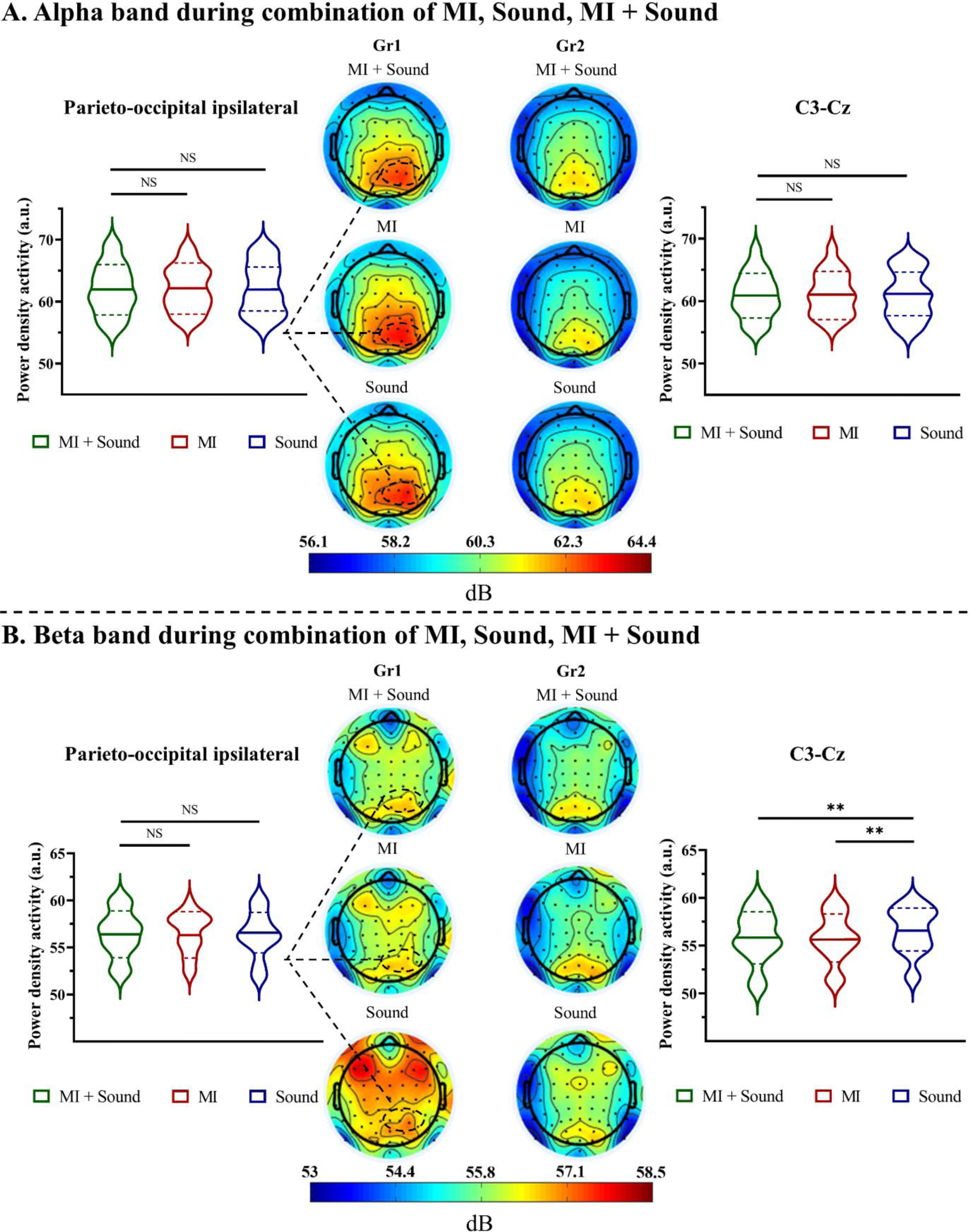
Alpha **(A)** and beta **(B)** band during MI, Sound, and MI + Sound for Group 1 and Group 2. The middle figure represents scalp topography of power density activity for the alpha (8–12 Hz) and beta (13–20 Hz) power during MI, Sound, and MI + Sound. For the right and left figures, violin plots represent power density activity. Thick and thin horizontal lines mark the mean and SD, respectively. ** = *p* < .01.

## 3. Discussion

The present study provides several notable findings related to the influence of MI and low-frequency sound listening on brain oscillations. First, our data not only reaffirm previous research findings but also extend our understanding by revealing diverse modulations within the alpha and beta frequency bands during MI and listening to low-frequency sounds. Nevertheless, these modulations exhibit substantial interindividual variability, particularly within the sensorimotor areas and parieto-occipital regions. Second, the combination of MI and low-frequency sound listening did not exhibit additional alpha or beta modulation compared to these processes in isolation.

Regarding the first point, a substantial body of research consistently highlights modulations in alpha and beta bands, often manifested by an event-related desynchronization or a power decrease in sensorimotor regions, accompanied by a concurrent increase in ipsilateral areas, particularly parieto-occipital regions (Brinkman et al., 2014; de Lange et al., 2008; Di Nota et al., 2017; McFarland et al., 2000; Pfurtscheller, 2000; Pfurtscheller & Neuper, 1997; Pfurtscheller at al., 1997; Putzolu et al., 2022). However, our findings point out a significant interindividual variability (Höller et al., 2013; Wriessnegger et al., 2020), demonstrating the diverse brain responses observed among our participants.

In particular, group 2 (comprising 8 individuals) supported the decrease in alpha and beta bands power within sensorimotor areas during MI, which closely mirrors the brain activity observed during actual movement (McFarland et al., 2000; Pfurtscheller, 2000; Toro et al., 1994). This reduction is thought to correspond to a heightened level of cortical activity, reflecting activated neural clusters ready or primed for sensory, motor, or cognitive processing. Conversely, group 1 (comprising 12 individuals) displayed increased alpha power within both sensorimotor and ipsilateral medial parieto-occipital regions. This increased power might be interpreted as an active inhibition or a disengagement of these cortical areas. Hence, we observed a wide-ranging variability phenomenon, which could be potentially attributed to disparities in kinaesthetic imagery capabilities (Toriyama et al., 2018).

Secondly, to the best of our knowledge, the present research marks an innovative effort in exploring brain oscillations while individuals are engaged in listening to low-frequency sounds. As observed for MI, our findings revealed a diverse range of modulations in both alpha and beta band power, depending on the cortical regions considered and the interindividual variability. More specifically, group 2 exhibited a selective decrease in alpha band power within sensorimotor areas. This pattern strongly suggests heightened sensorimotor activity during the process of listening to low-frequency sounds, akin to the patterns observed during MI. Furthermore, group 1 displayed a noteworthy increase of alpha and beta band power within ipsilateral medial parieto-occipital regions, suggesting an active inhibition or a disengagement of these cortical areas. These observations bear a remarkable resemblance to our MI findings, and corroborate our positive correlations between MI and Sound, further emphasizing the hypothesis that combining both tasks could elicit increased modulation of alpha and beta band power.

Nonetheless, it is noteworthy that the combination of MI and low-frequency sound listening did not yield any further increase or decrease in alpha or beta band power for the two investigated brain regions. This finding is rather surprising in light of recent research suggesting potential benefits of low-frequency sound listening for improving motor rehabilitation (Chatain et al., 2018; Naghdi et al., 2015). This may be explained by an interference effect, indicating that when MI and low-frequency sound listening are performed simultaneously, they could lead to a significant interference, characterized by overlapped and concurrent brain activations (Klingberg & Roland, 1997). Therefore, further potentiation is barely noticeable with the combination of both at the same time. In addition to our cortical findings, some studies propose that enhanced motor performance or rehabilitation following auditory stimulation may be attributed to an audiospinal facilitation phenomenon, characterized as an enhanced excitability in motoneurons due to a reduction in presynaptic inhibition (Fernandez-Del-Olmo et al., 2014; Rossignol & Jones, 1976).

Our main findings must be considered in the light of the study limitations. First, our decision to employ clustering analysis was deliberate and deemed appropriate for our research objectives. However, this methodological choice resulted in a constrained number of participants allocated to each group, reducing the statistical power of our analysis. However, despite this limitation, the effect sizes (Cohen’s *d* or partial eta squared) for MI and low-frequency sound listening tend towards large effects, thus confirming the validity of our findings. A second limitation is the subjective measurements of the imagery ability using the revised version of Movement Imagery Questionnaire. Indeed, we specified above that the interindividual variability could be potentially attributed to disparities in kinaesthetic imagery capabilities (Toriyama et al., 2018). However, we cannot ensure that our interindividual differences stem for different kinaesthetic MI abilities as we did not separately assess the various modalities of MI. In addition, while subjective assessments serve as an index of imaging ability, integrating an objective measure as the amplitude of motor evoked potentials through transcranial magnetic stimulation (TMS) would have bolstered our study. Indeed, some researchers have demonstrated that the modulation of these responses during MI correlates with the participants’ imagery ability (Lebon et al., 2012). Nevertheless, recent findings have tempered this concern by indicating an absence of correlation between responses recorded via EEG and TMS (Vasilyev et al., 2017).

To conclude, our results shed significant insights into the effects of MI and low-frequency sound listening on brain oscillations and contribute to the idea that individuals cannot be uniformly categorized, as observing group-level effects may not always reflect the underlying reality. Second, our results indicate that although MI and low-frequency sound listening elicit similar (albeit not identical) brain oscillations, the combination of both does not automatically produce additional modulations within sensorimotor or parieto-occipital regions. Further investigations are recommended to explore the potential influence of timing in the interplay of two interventions, specifically, whether an additional modulation of brain activity becomes apparent when both interventions are administered sequentially. Such research could provide valuable insights into the temporal dynamics of these interventions and their combined effects on neurophysiological modulations.

## Supporting information

Supplementary section

## Data availability statement

All data from this study are available at: https://osf.io/jnvkp/?view_only=58e58960b7424581bf1d02c8d7d0a10b

## Author contributions

Experiment design: CML, FL, and PCB; Data collection: WD and PCB; Statistical analysis: WD, PCB and JT; Manuscript preparation: WD, CML, FL; Manuscript review: PCB, JT and DAT.

