## Supplementary section for "Exploring dynamic brain oscillations in motor imagery and low-frequency sound"

### Supplementary materials

S1. $EMG_{rms}$ activity (mean ± SD) in μV recorded for each condition

S2. Silhouette scores and number of clusters

S3. Comparison of the rest between the two groups

S1. $EMG_{rms}$ activity (mean ± SD) in µV recorded for each condition

**Table S1**. $EMG_{rms}$ activity (mean ± SD) in µV recorded for each condition. We analyzed the $EMG_{rms}$ during all tasks to ensure that alpha band activity was not influenced by muscle contractions. We did not observe any difference between baseline and the other conditions [Friedman ANOVA FDS: $r(3) = 0.039$, $p = 0.166$; Friedman ANOVA EDS: $r(3) = -0.024$, $p = 0.643$]. This result demonstrated that alpha and beta power modulations were not influenced by muscular preactivity.

| | **Rest** | **MI** | **Sound** | **MI + Sound** |
|---|---|---|---|---|
| **FDS Mean ± SD (µV)** | 2.58 ± 1.62 | 2.82 ± 1.55 | 2.54 ± 1.52 | 2.65 ± 1.76 |
| **EDS Mean ± SD (µV)** | 4.64 ± 3.17 | 4.36 ± 2.34 | 4.22 ± 2.77 | 4.91 ± 2.62 |

S2. Silhouette scores and number of clusters.

**Table S2.** Silhouette scores and number of clusters

| **Number of clusters** | **Silhouette Scores** |
|---|---|
| 2 | 0.667 |
| 3 | 0.572 |
| 4 | 0.548 |

S3. Comparison of the rest between the two groups

Repeated measures ANOVA revealed no main effect of Group, $F(1, 18) = 0.004$, $p = .952$, $\eta_p^2 = 0.0002$, nor any interaction between Group, Bands and ROIs, $F(1, 18) = 0.074$, $p = .789$, $\eta_p^2 = 0.004$, indicating no significant difference in power amplitudes, irrespective of band of interest (alpha and beta) or regions of interest (sensorimotor or parieto-occipital).
